# Struct2SeQ: RNA inverse folding with Deep Q-Learning

**DOI:** 10.64898/2026.01.16.700031

**Authors:** Shujun He, Qing Sun

## Abstract

RNA molecules play critical roles in biology and therapeutics, with their function intimately tied to their secondary structure. Designing RNA sequences that reliably fold into desired secondary structures, especially those with complex pseudoknots, remains a fundamental challenge. Here, we present Struct2SeQ, a reinforcement learning framework that leverages deep Q-learning to generate RNA sequences conditioned on target secondary structures and SHAPE reactivity constraints. By formulating RNA design as a sequential decision-making process, our model learns to construct sequences that not only fold into the intended structures but also exhibit experimentally consistent SHAPE profiles. Evaluated with experimental data from the OpenKnot 240mer pseudoknot design challenges, Struct2SeQ significantly outperformed humans and other automated design methods while generating diverse solutions that explore a broader sequence space compared to human players. The incorporation of SHAPE-informed rewards further enhances the chemical validity of generated sequences, as evidenced by improved OpenKnot scores. Our results demonstrate the potential of reinforcement learning for RNA design tasks, opening avenues for engineering RNAs with complex structures and functions.

## 1 Introduction

RNA molecules play a central role in numerous biological processes, including gene regulation, catalysis, molecular recognition, and protein translation [1]. Unlike DNA, the function of RNA is intimately tied not only to its sequence, but also to its three-dimensional shape which is mostly driven by secondary structure (i.e. base pairing). Designing RNA sequences that reliably fold into desired secondary structures is a fundamental challenge in RNA biology [2]. Solving this fundamental challenge will allow a myriad of applications, including engineering of riboswitches and aptamers with specific ligand-binding properties [3], the development of guide RNAs for CRISPR systems that satisfy structural constraints [4], the creation of mRNA therapeutics with optimized stability and translation efficiency [5, 6, 7, 8], and the construction of RNA biosensors and synthetic gene control circuits [9]. Recently, foundation models have been trained on massive amounts of chemical mapping data from both natural and synthetic sources [10]. However, there is a finite amount of natural RNAs that can be used for training and previous synthetic RNAs were crowd sourced. Therefore, both are not scalable enough for million and billion scale RNA sequence generation. As a result, it is important to have a scalable framework to generate RNA sequences with interesting structures.

Despite the potential applications, RNA design remains difficult due to the many-to-one mapping from sequence to structure, the sparsity of sequence-structure pairs, and the limitations of folding energy models. Furthermore, existing design tools often rely on heuristic search or energy minimization and cannot flexibly accommodate additional chemical constraints such as SHAPE reactivity profiles or experimental feedback.

To address these challenges, we propose a reinforcement learning-based framework that directly learns to generate RNA sequences from target structures. By leveraging deep Q-learning and dense, chemically informed rewards derived from accurate neural predictors of folding and chemical reactivity (RibonanzaNet-SS and RibonanzaNet), our model enables structure-conditioned generation of RNA sequences that conform to both structural and chemical constraints.

## 2 Methods

### 2.1 Environment

We use two versions of RibonanzaNet as the environment that provides reward signals to the reinforcement learning agent. RibonanzaNet is a deep neural network designed for RNA structure-function prediction, capable of mapping a given RNA sequence to either a SHAPE reactivity profile or a base-pairing pattern.

#### Structure-Predictive Environment

The first version of RibonanzaNet is RibonanzaNet-SS, which is fine-tuned to predict secondary structure as a base-pair map *ŷ*^struct^ = {(*i, j*)}, where each predicted pair indicates that nucleotides at positions *i* and *j* are likely to form a canonical base pair (e.g., A–U, G–C, G–U). This version serves as the primary structural feedback mechanism, allowing us to determine whether the generated RNA folds into the desired target structure *in silico*. The output is compared against the ground truth structure to compute per-position binary rewards based on pairing correctness.

#### SHAPE-Predictive Environment

The second version of RibonanzaNet is the base model pretrained on experimental chemical mapping data (e.g., SHAPE-Seq) and outputs a predicted SHAPE reactivity profile *ŷ*^shape^ ∈ ℝ^*L*^, where higher values indicate greater nucleotide flexibility or solvent accessibility (typically corresponding to unpaired positions). These predicted reactivities are used to impose biochemical constraints on the generated sequences.

#### Reward Integration

Both RibonanzaNet models serve as efficient proxies for traditional RNA folding simulations. The structure-predictive model enables position-level rewards for correct base-pairing, while the SHAPE-predictive model enables constraint-based filtering using thresholds on predicted reactivities. Together, they define a dense and chemically grounded reward function of the form:

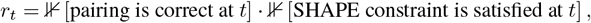

which is computed for every position in the sequence and summed across the sequence to obtain the total reward.

#### Chemical Plausibility

By relying on neural predictors trained on experimental data, the environment provides reward signals that are more closely aligned with real-world RNA behavior than purely algorithmic folding tools. This also allows for generalization to structural motifs or chemical profiles not seen during training.

Overall, this dual-environment setup enables Struct2SeQ to generate sequences that are not only structurally accurate but also chemically plausible according to experimentally derived constraints.

### 2.2 Reinforcement learning

The main idea of deep Q-learning is to train a neural network to predict the the Q-value[11], a weighted sum of immediate reward and future action values; the weighted sum represents the neural network’s belief about how good an action is given a state for all future steps combined. Mathematically, the Q-value *Q*(*s, a*) represents the expected cumulative reward for taking action *a* in state *s*, and following the optimal policy thereafter. The core update is based on the Bellman equation:

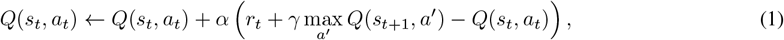

where:

- *α* is the learning rate,
- *γ* ∈ [0, 1) is the discount factor,
- *r*_*t*_ is the reward at time *t*,
- *a*^*′*^ denotes the next possible action at *s*_*t*+1_.

In deep Q-learning, we approximate *Q*(*s, a*) using a neural network *Q*_*θ*_(*s, a*) with parameters *θ*. The model is trained to minimize the temporal difference loss:

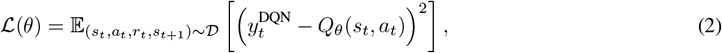

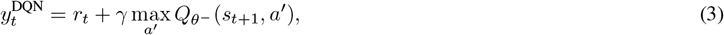

where 𝒟 is the replay buffer and 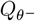 is a target network whose parameters are periodically synchronized with *θ*. This synchronization helps stabilize training by decoupling the target values from the potentially rapidly changing network.

### 2.3 Sequence to Sequence Modeling with Deeq Q learning

The task of RNA secondary structure design involves generating an RNA sequence that folds into the target secondary structure. This target is often specified using dot-bracket notation, which encodes base-pairing interactions and unpaired regions in a compact symbolic form. From a machine learning perspective, this can be framed as a sequence-to-sequence (seq2seq) problem, where the input is a dot bracket structural representation and the output is a compatible RNA sequence.

Neural machine translation sequence-to-sequence (seq2seq) models learn a direct mapping between input and output sequences using supervised learning on large datasets of paired examples [12]. Because standard seq2seq models optimize token-level likelihood, they do not explicitly account for the global structural constraints of RNA folding. Further, data of RNAs with known secondary structures is scarce. As a result, directly training a conventional sequence-to-sequence model to generate sequences from target secondary structures with supervised learning is not likely to work well.

To better align the model’s objective with the downstream structural constraints, we adopt a reinforcement learning (RL) framework, specifically deep Q-learning. In this setting, the model acts as an agent that incrementally constructs a sequence, making a discrete decision (nucleotide selection) at each position. After generating a full sequence, the agent receives a scalar reward at each position based on if that position is correctly paired or unpaired and if that position meets SHAPE constraints as defined by OpenKnot score.

We formulate the generation process as a Markov Decision Process (MDP) as follows:

- **State** *s*_*t*_: the current partially generated sequence up to position *t*, along with the full target structure in dot-bracket notation. Optionally, additional features like pair indices or local structure context can be included.
- **Action** *a*_*t*_ ∈ {A, U, G, C} : the nucleotide selected at position *t*. For positions that are base-paired in the target structure, the selected right nucleotide must be compatible with the canonical base-pairing rules (e.g., A–U, G–C, G–U) relative to its pairing partner.
- **Transition**: upon taking action *a*_*t*_, the nucleotide is appended to the sequence, and the environment moves to the next position *t* + 1.
- **Reward** *r*_*t*_ ∈ ℝ: a local reward given at each position, defined based on if the nucleotide placed at *t* is correctly paired or unpaired and if that position meets SHAPE constraints as defined by OpenKnot score.
- **Q-value** *Q*_*θ*_(*s*_*t*_, *a*_*t*_): a neural network parameterized by *θ* that predicts the expected cumulative reward for choosing action *a*_*t*_ in state *s*_*t*_, and following the learned policy thereafter.

This framework enables the model to learn a generation policy that respects the structural requirements of RNA folding, guided directly by chemically meaningful rewards rather than likelihood-based supervision.

### 2.4 Reward Design

A key component of our reinforcement learning framework is the design of a structurally and chemically grounded reward function. Rather than relying solely on predefined rules or physics-based folding simulations, we leverage a neural network predictor to estimate secondary structure and SHAPE reactivity profiles from candidate sequences. This allows us to define position-wise rewards that reflect the agreement between predicted and target structure-specific constraints, while incorporating SHAPE data predicted *in silico*.

To guide the RNA sequence generation process toward chemically valid and structurally accurate solutions, we define a dense, position-wise reward function based on two discrete criteria: (1) correct base-pairing status, and (2) compliance with SHAPE reactivity constraints.

#### Secondary Structure Representation

The target secondary structure is provided as a set of base-pair mappings *y*^struct^ = {(*i, j*)}, where position *i* is base-paired with *j*. Unpaired positions are those not included in any mapping. The predicted structure *ŷ*^struct^ is represented in the same way based on predictions from RibonanzaNet.

#### SHAPE Constraints

Each position *t* also has an associated SHAPE constraint, based on predictions from a neural RibonanzaNet. We use a binary rule-based threshold:

- Positions predicted to be ^**^unpaired^**^ should have SHAPE reactivity above a given threshold (e.g., *>* 0.5),
- Positions predicted to be ^**^paired^**^ should have SHAPE reactivity below a different threshold (e.g., *<* 0.25).

The predicted SHAPE profile *ŷ*^shape^ ∈ ℝ ^*L*^is obtained from a neural network applied to the generated sequence.

#### Reward Function

At each position *t*, the reward *r*_*t*_ is defined as a product of two binary terms:

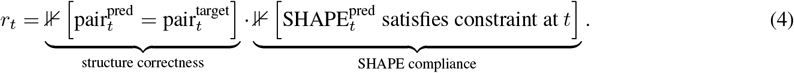

This yields:

- *r*_*t*_ = 1 if the pairing status is correct and the SHAPE constraint is satisfied,
- *r*_*t*_ = 0 otherwise.

By enforcing base-pairing fidelity as a prerequisite and incorporating SHAPE compliance only when structure is correct, this reward encourages the generation of sequences that not only fold into the intended secondary structure, but also exhibit accessibility patterns consistent with experimentally observed reactivity. This reward function provides dense supervision throughout sequence generation, enabling effective credit assignment at each step. It encourages the model to produce sequences that result in both globally accurate folding patterns and SHAPE profiles consistent with secondary structure constraints.

In practice, we trained two variants of this model with different reward functions: Struct2SeQ, which uses only the structure correctness term, and Struct2SeQ-SHAPE, which incorporates both structure correctness and SHAPE compliance. We restart training from the Struct2SeQ checkpoint to obtain Struct2SeQ-SHAPE. This allows us to assess the impact of SHAPE-informed rewards on sequence design quality.

### 2.5 Model architecture

Our model for RNA sequence generation conditioned on secondary structure combines convolutional, transformer-based [12], long short term memory (LSTM) [13], and graph convolutional modules [14] to capture both sequential and structural dependencies inherent in RNA folding. The architecture follows a sequence-to-sequence encoder-decoder paradigm, consisting of a dot-bracket-aware encoder and a structure-guided decoder.

#### Overview

Given an input target secondary structure represented in dot-bracket notation, the model encodes the structure into contextual embeddings using a transformer based encoder. The decoder then autoregressively generates an RNA sequence, producing one nucleotide at a time. The model is designed to integrate both sequential information (via transformers and convolutions) and structural information (via graph convolutions and contact maps).

#### Encoder

The encoder processes the input secondary structure using several components:

- **Embedding Layer:** The dot-bracket tokens are embedded into a continuous space using a learned embedding table.
- **Positional Encoding:** A learned LSTM-based positional encoder is used to capture the relative ordering of tokens.
- **Convolutional Feature Extraction:** A 1D convolutional layer captures local structural motifs over the embedding space.
- **Graph Convolution:** A structure-aware graph convolutional network (GCN) integrates pairwise contact information via the input contact matrix. This helps the model to reason about long-range base-pairing interactions beyond local neighborhoods.
- **Transformer Encoder:** Finally, a stack of Transformer encoder layers aggregates global context via multi-head self-attention.

#### Decoder

The decoder is a causal transformer that generates the RNA sequence token-by-token:

- **Token Embedding:** The decoder receives the partial output sequence as input and embeds it via a learned embedding table.
- **Paired Encoding Injection:** A binary paired/unpaired encoding, derived from the structure, is embedded and added to the decoder input to explicitly condition on pairing status.
- **LSTM Positional Encoding:** Similar to the encoder, positional information is added using a learned LSTM-based encoding.
- **Causal Convolution:** A causal convolutional layer ensures that each position only attends to past tokens, preserving autoregressive decoding.
- **Causal Transformer Decoder:** A custom Transformer decoder supports key-value caching for efficient step-wise generation during inference. It includes masked self-attention, encoder-decoder cross-attention, and position-wise feedforward layers.
- **Output Projection:** The final hidden states are projected into the vocabulary space (excluding the padding token), yielding Q values over the four nucleotide classes.

### 2.6 Playing and Training

We adopt a variant of Deep Q-Learning with experience replay for sequence to sequence modeling, which we refer to as **twice-shifted action value training** (fig. 2). This approach arises naturally in the autoregressive setting, where both the decoder input and the target Q-values are temporally shifted. There are two main components during training, playing against the environment and training the agent using replay experience of structure sequence pairs with gradient descent. We summarize these 2 components in algorithm 1 and algorithm 2.

**Figure 1:**
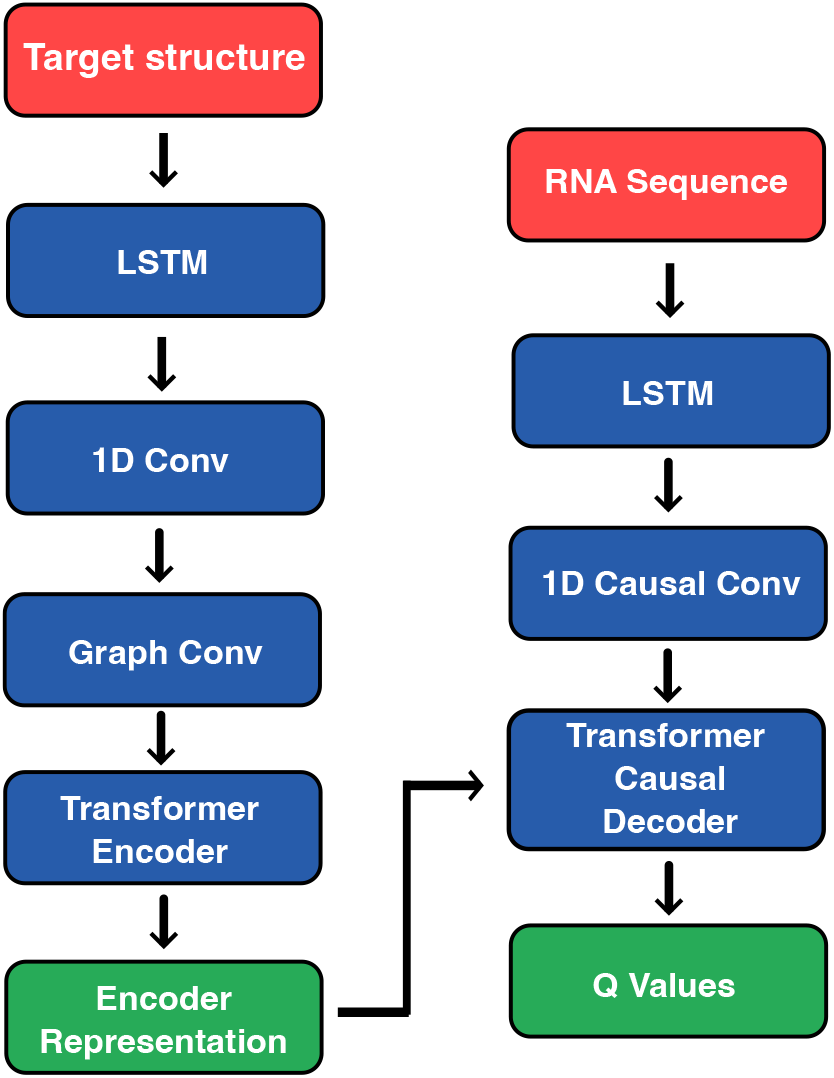
Neural network architecture of Struct2SeQ. The model consists of a dot-bracket-aware encoder and a structure-guided decoder, integrating convolutional, transformer, and graph convolutional modules to capture both sequential and structural dependencies.

**Figure 2:**
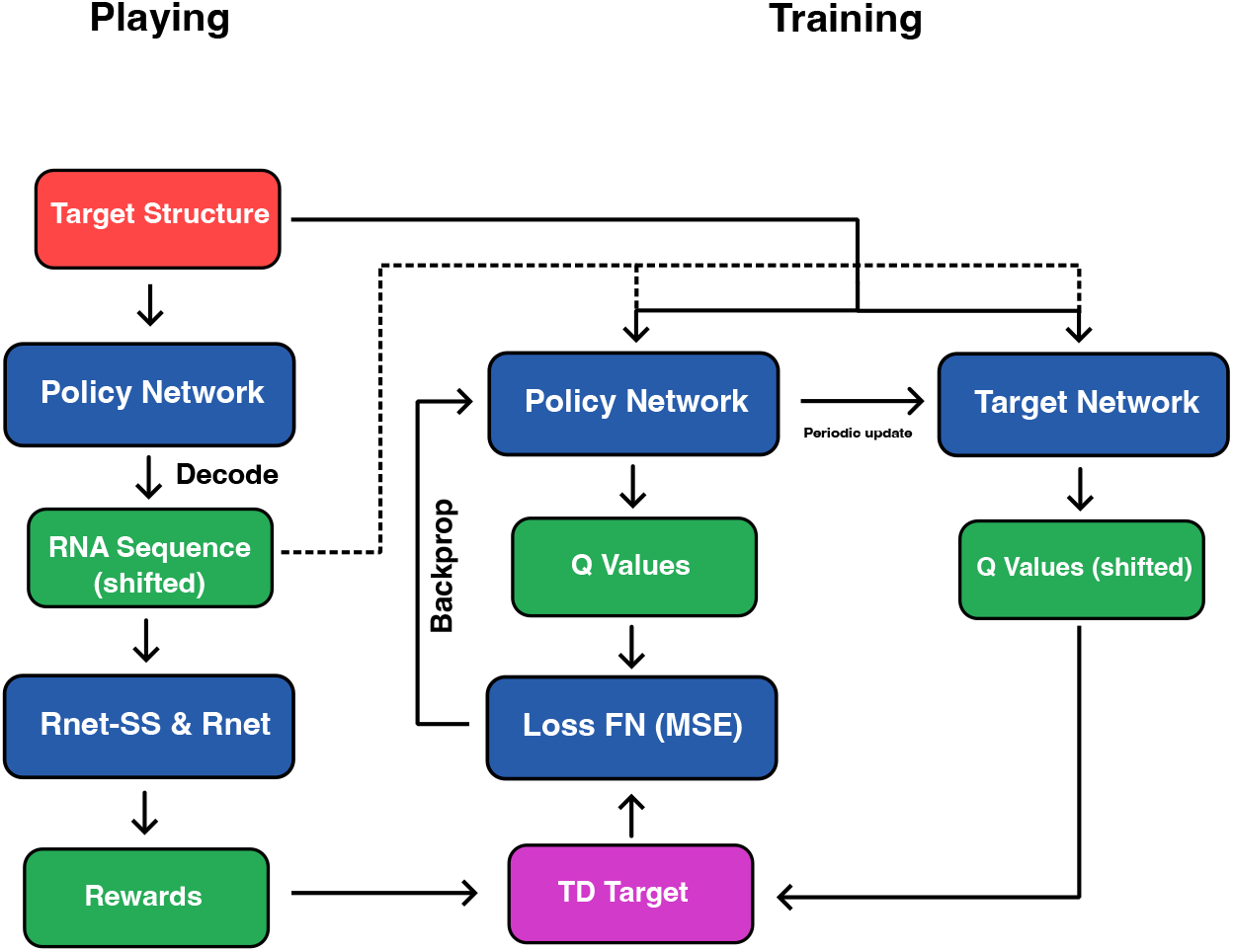
Playing and twice-shifted action value training for sequence-to-sequence deep Q-learning. Both the decoder input and the target Q-values are temporally shifted to enable autoregressive generation while propagating reward signals across time.

**Figure 3:**
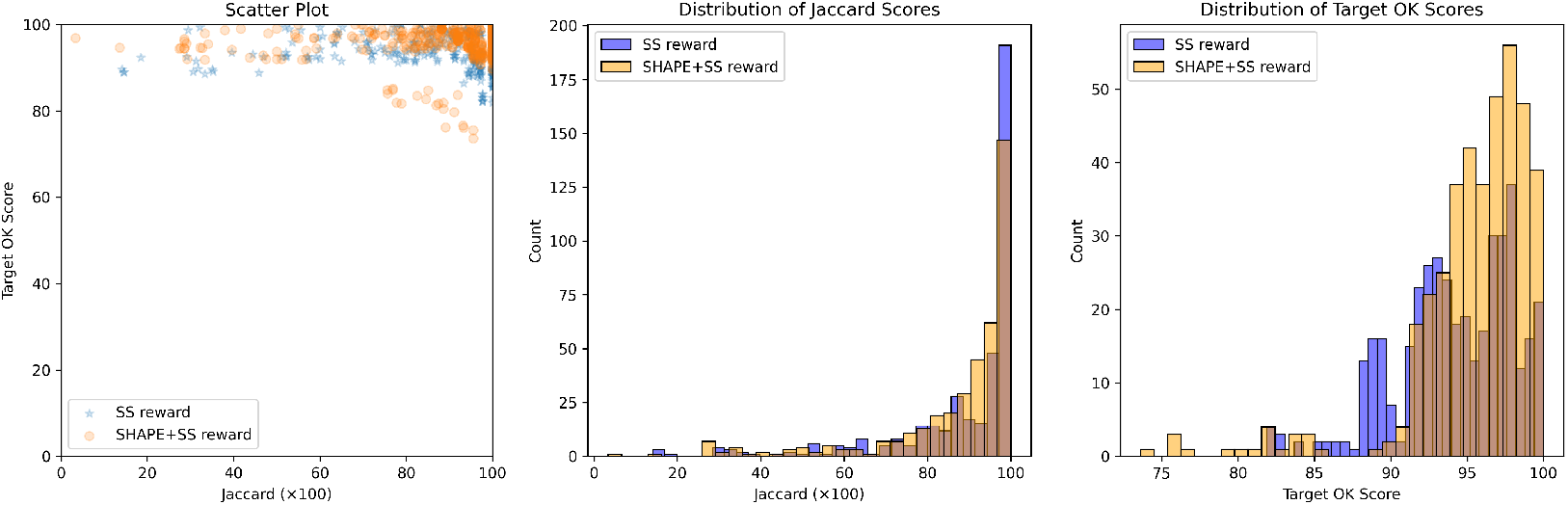
Performance of Struct2SeQ and Struct2SeQ-SHAPE on the OpenKnot 7 challenge. Left: Jaccard of base-pairs score for generated sequences vs scatter plot of Target OpenKnot score. Middle: histogram of Jaccard of base-pairs scores for generated sequences. Right: histogram of Target OpenKnot scores for generated sequences

#### Twice-shifted Action Value Training

At each training step, the model receives a sequence of structure tokens (dot-bracket notation) and a ground truth RNA sequence. The decoder is trained to predict Q-values for each possible nucleotide at each position, aiming to maximize the expected cumulative reward associated with generating a correct and SHAPE-compliant RNA sequence. To enable autoregressive generation, the input to the decoder is the RNA sequence shifted by one position to the right, with a start token appended to the front. This ensures that the prediction at position *t* is conditioned only on the tokens *<* 1, …, *t* − 1 *>*, satisfying causality. Simultaneously, the target Q-values used for temporal-difference updates are also shifted: the maximum Q-value at position *t* + 1 is used to compute the target for position *t*. As a result, both the input sequence to the decoder and the temporal target are shifted and hence we call this training process **twice-shifted action value training**.

While technically the reward function cannot be computed until the entire sequence is generated, we can still use the reward signal from the full sequence to assign position-wise rewards during training, since we have generated the full sequence during playing. When training with many examples, this can be interpreted as learning the expected future reward at each position, which is exactly what the Q-value represents.

#### Loss Function

The training loss is the mean squared error between the predicted Q-value for the selected action and the TD target:

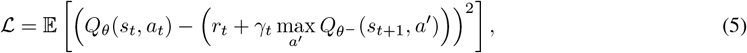

where:

- *Q*_*θ*_(*s*_*t*_, *a*_*t*_): predicted Q-value for the selected action *a*_*t*_,
- 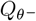: target network (frozen copy of policy),
- *γ*_*t*_: linearly decayed discount factor across the sequence,
- *r*_*t*_: scalar reward at position *t*.

#### Reward Reweighting

Generally, it is probably much more difficult to go from near perfect sequences (e.g., 95% correct) to perfect sequences (100% correct) than from poor sequences (e.g., 10% correct) to near perfect sequences. Therefore, if the reward function has linear scaling, there may not be sufficient gradient to push the model to improve near-perfect sequences to perfection, so naturally we scale up the reward for better sequences exponentially to create enough gradient, where the reward vector is reweighted by an exponential scaling factor based on its sequence-level mean:

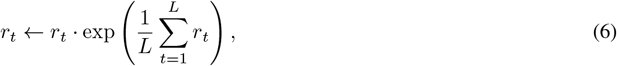

which boosts higher-scoring examples.

#### Position-Dependent Discounting

Unlike standard deep Q-learning, which uses a fixed discount factor *γ*, we apply a linearly decayed discount factor along the sequence dimension to encourage the model to think more about future values at early positions, which have greater influence on global structure formation. The discount schedule is defined as:

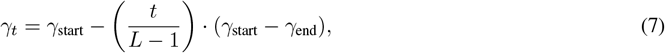

where *L* is the sequence length, and *γ*_start_ *> γ*_end_. This formulation encourages the model to focus more on early decoding decisions that have disproportionately high impact on downstream generation.

#### Optimization Details

Training is performed with mixed precision with Adam. Multi-GPU and multi-node training are supported via the `Accelerate` framework. Gradients are clipped to prevent instability, and a cosine learning rate scheduler is applied. The target network remains fixed during each epoch and is updated periodically during training.

#### Masking

A square causal mask is applied to enforce the autoregressive constraint.

##### Algorithm 1

Play Episode and Collect Rewards

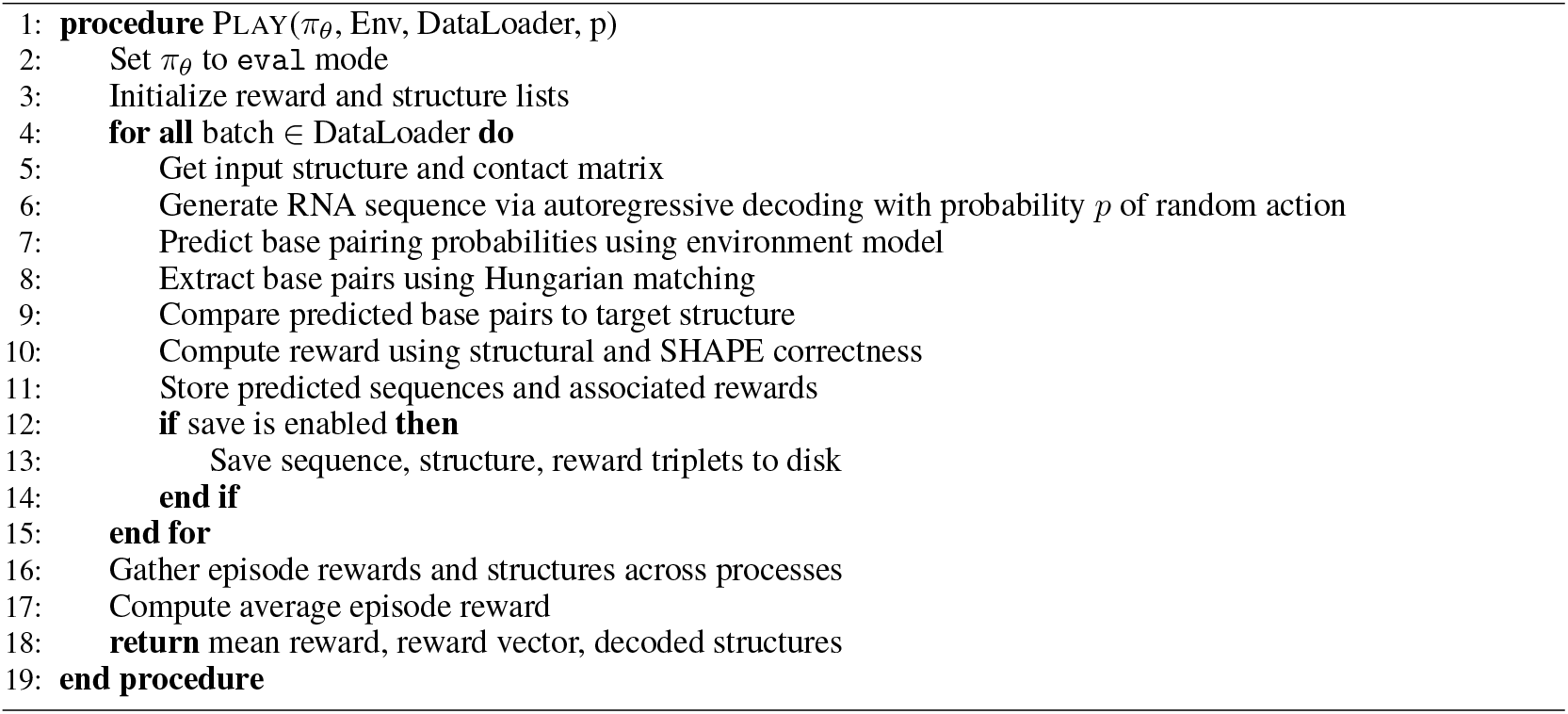

##### Algorithm 2

Twice-Shifted Action Value Training

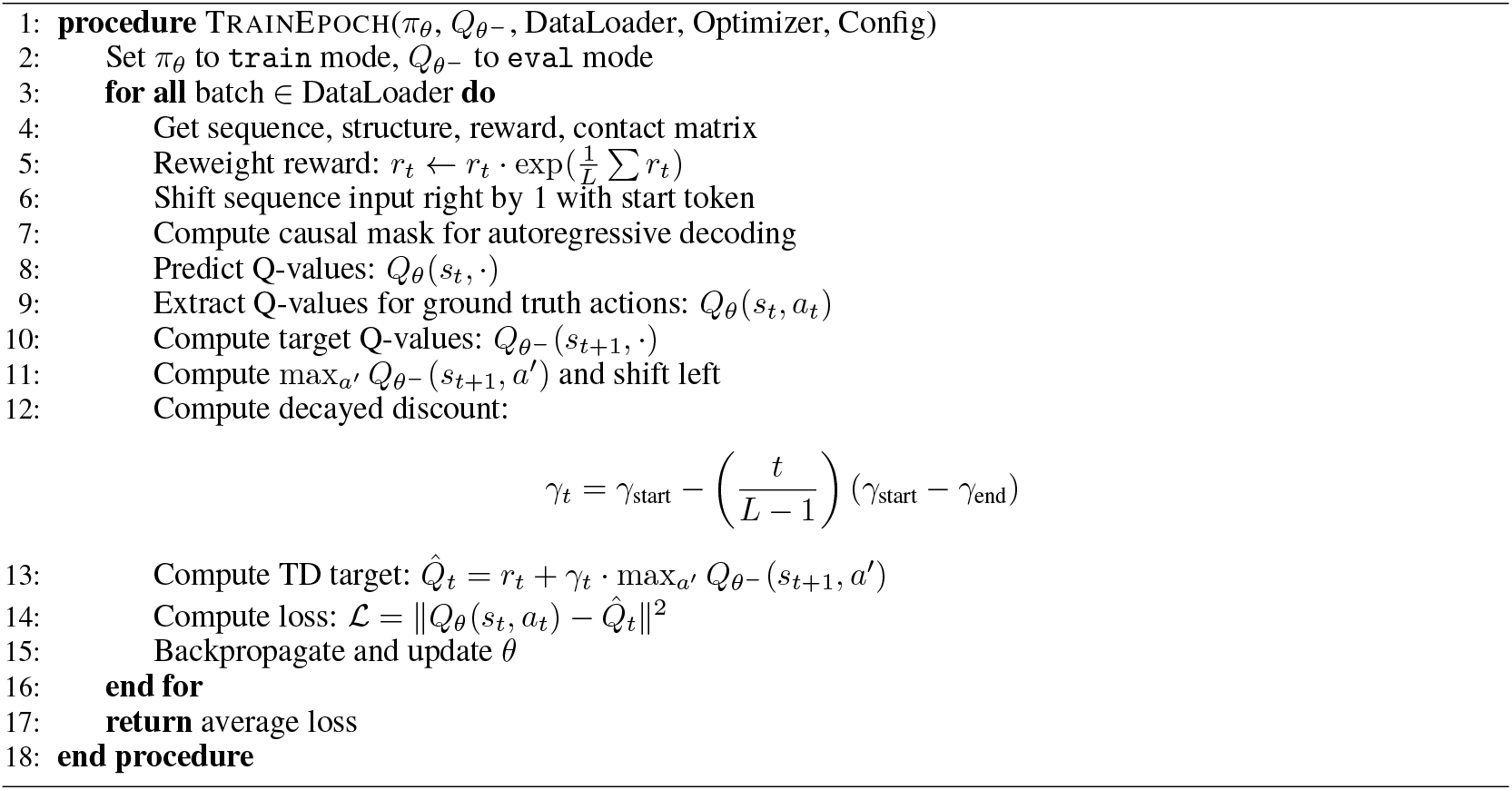

### 2.7 Inference

At inference time, Struct2SeQ operates as an autoregressive generative model that decodes RNA sequences conditioned on a target secondary structure. The model outputs action-value estimates (Q-values) for each nucleotide choice at each position, and uses these estimates to guide sequence generation under structural constraints.

#### Autoregressive Decoding

The model generates sequences one nucleotide at a time, conditioning each prediction on the previously generated tokens and the encoded dot-bracket structure. At each time step *t*, the model predicts a Q-value distribution *Q*(*s*_*t*_, ·) over the four nucleotides {A, U, G, C}. Inference strategies determine how these Q-values are used to select the next nucleotide.

#### Beam Search

One decoding strategy is beam search. At each step:

- The top-*k* sequences with the highest cumulative Q-values (i.e., sum of Q-values across all positions so far) are retained.
- For each of the *k* sequences, the model considers all possible next actions, selects the top-*k* expansions, and continues.

This approach offers good average performance and generates sequences that reliably fold into the desired structure. However, it tends to produce relatively homogeneous outputs, especially when the Q-values are sharply peaked. We note that while beam search is typically used with summed log-probabilities which can be interpreted as conditional likelihoods, here we use cumulative Q-values which do not have a direct probabilistic interpretation. Nevertheless, selecting sequences with the highest cumulative Q-values effectively identifies those expected to yield the best structural fidelity according to the learned Q-value.

#### Random Search

To enhance diversity, we optionally employ a random search strategy:

- With some probability, the next nucleotide is sampled from a softmax-transformed Q-distribution or chosen uniformly over valid nucleotides (e.g., allowed base pairs).
- This encourages exploration and allows for the discovery of highly divergent but still valid solutions.

#### Rescue Strategy

For cases where generated sequences almost match the target (e.g., one or two incorrect base pairs), we apply a local repair procedure:

- Identify mismatch positions,
- Enumerate all valid nucleotide combinations over those positions,
- Re-evaluate structural fidelity and select the corrected sequence with the best folding match.

While not scalable for large batch inference, this is useful in design settings where near-miss sequences can be easily corrected to exact solutions.

#### Screening

Generated sequences are filtered using post-hoc evaluation metrics:

- **Jaccard of base-pairs score**: intersection over union of predicted and target base-pair sets, which measures agreement between predicted and target structures,
- **OpenKnot score**: Quantifies how well an RNA design is supported by experimental SHAPE data and is compatible with target base pair formation with special consideration for pseudoknots. The score (0–100) is the mean of (i) the *Eterna Classic Score*, which penalizes predicted paired residues with SHAPE *>* 0.5 and predicted unpaired residues with SHAPE *<* 0.125, assuming SHAPE values normalized to a 90th–percentile of 1.0; and (ii) the *Crossed Pair Quality Score*, computed identically but restricted to residues involved in crossing base pairs (with singlets removed and flanking-region partners downweighted). We follow the implementation from https://github.com/eternagame/OpenKnotScorePipeline.

These screening metrics enable selection of high-quality candidates for downstream experimental validation or database inclusion.

Struct2SeQ supports both deterministic and stochastic inference, making it adaptable to diverse use cases: beam search for reliable outputs, random sampling for diverse solutions, and rescue for exact structural matches. Screening provides an additional layer of quality control to ensure that generated sequences satisfy both computational and experimental design criteria.

## 3 Results

### 3.1 OpenKnot Round 7 Pseudoknot design (100mer)

We participated in the OpenKnot 7 100mer challenge for designing RNA sequences that fold into pseudoknotted structures. For training we used 120 A6000 hrs (15hr on 8xA6000 node), and for inference, we set a compute budget where we generated 98000 sequences per target structure. We found that both Struct2SeQ and Struct2SeQ-SHAPE were able to generate sequences that match the target secondary structures at human-level performance while conforming to SHAPE constraints, although Struct2SeQ-SHAPE generated sequences with better OpenKnot scores while slightly underperforming in terms of number of puzzles solved (fig. 8). It is plausible that the SHAPE constraints narrow the sequence space and make it more difficult to find sequences that match the target secondary structure, but at the same time help generate sequences that are more chemically plausible. Overall, Struct2SeQ-SHAPE was able to generate sequences with better OpenKnot scores compared to Struct2SeQ, indicating that the SHAPE constraints help generate sequences that are more chemically plausible. However, we note that Eterna players were not provided the SHAPE constraints during the challenge, so we do not directly compare OpenKnot scores between Eterna players and Struct2SeQ-SHAPE. Overall, these results demonstrate the effectiveness of our reinforcement learning framework for RNA pseudoknot design, achieving human-level performance in a challenging design task.

Struct2seQ was able to solve 19/20 puzzles in terms of matching secondary structure (fig. 4), whereas Struct2SeQ-SHAPE was able to solve 18/20 puzzles (fig. 5). On the other hand, Eterna players were able to also solve 19/20 puzzles, so Struct2seQ was able to match human performance in this challenge when it comes to matching secondary structure. Strikingly, Struct2SeQ was also able to generate far more unique solutions compared to Eterna players, which simply cannot match the throughput of computer methods. While Eterna players typically have to start from the wild-type sequence and make mutations, which results in designs with small mutation distances (can we get data and add plot for this?), Struct2SeQ and Struct2SeQ-SHAPE were able to generate solutions de novo where designs are on average more than 50 mutations away from the wild-type sequence (fig. 6), indicating that our model is able to explore a much larger and distinct sequence space compared to human players.

**Figure 4:**
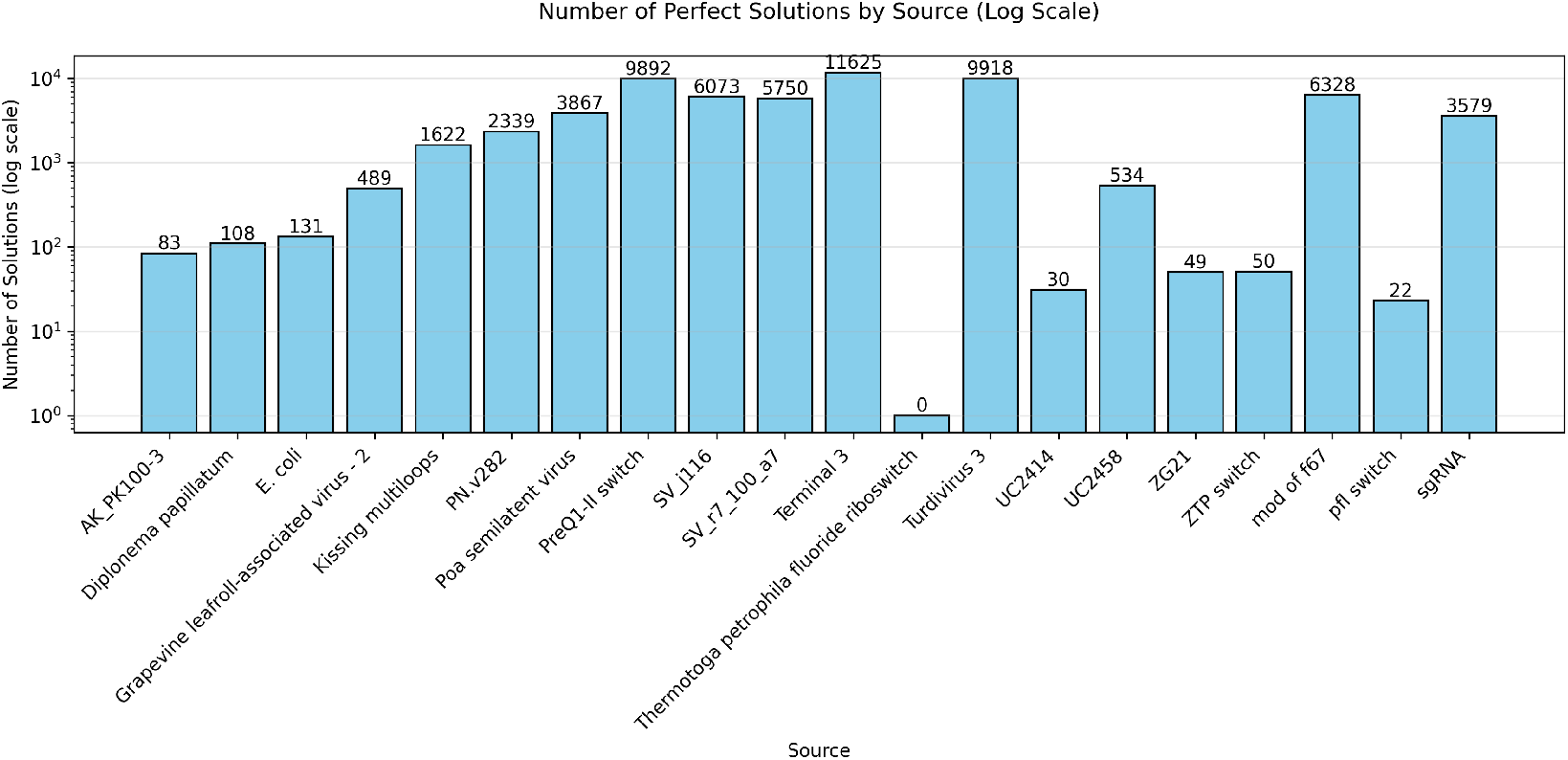
Number of solutions (matching base pairs) for each puzzle found by Struct2SeQ.

**Figure 5:**
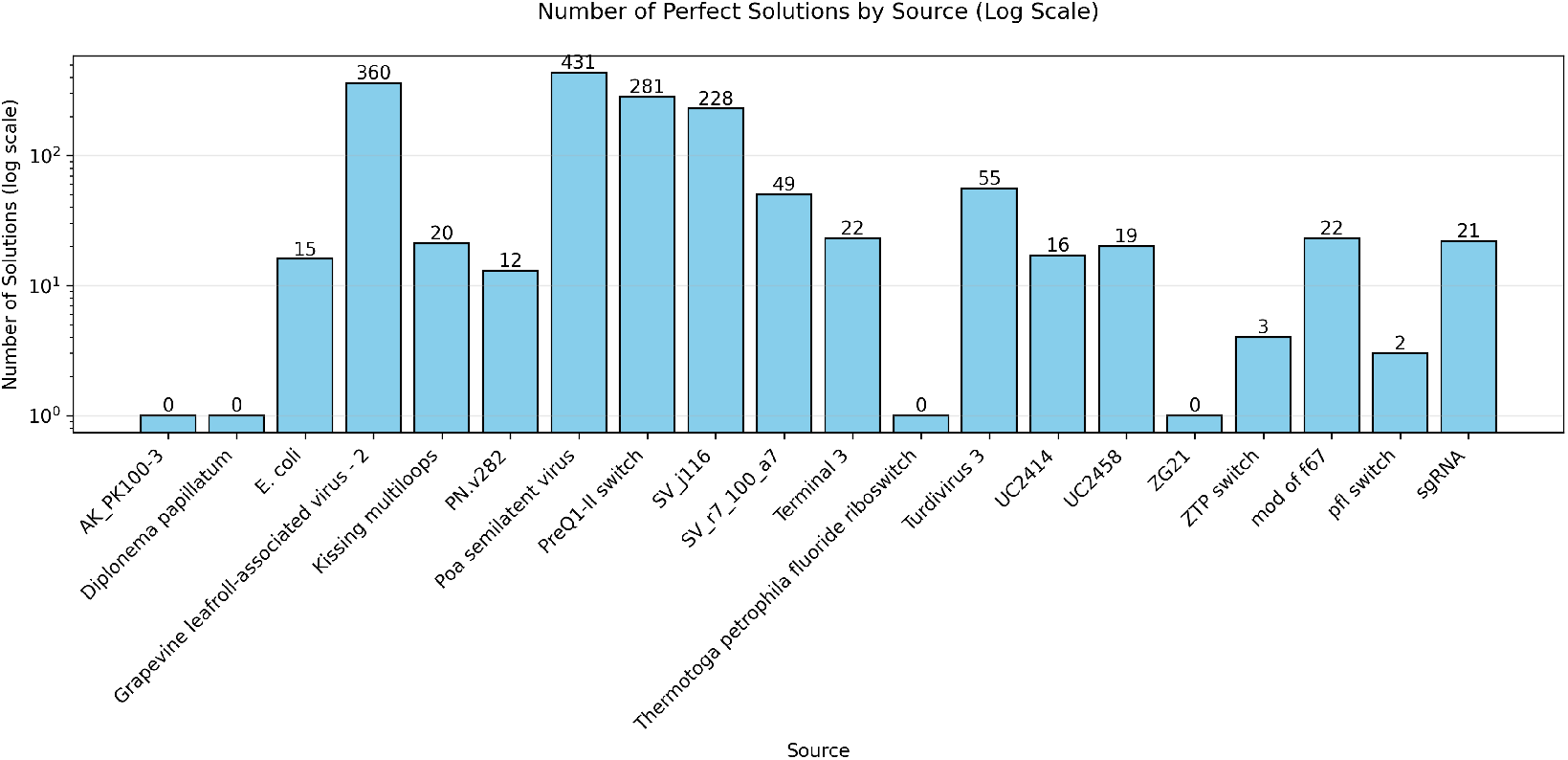
Number of solutions (matching base pairs) for each puzzle found by Struct2SeQ-SHAPE.

**Figure 6:**
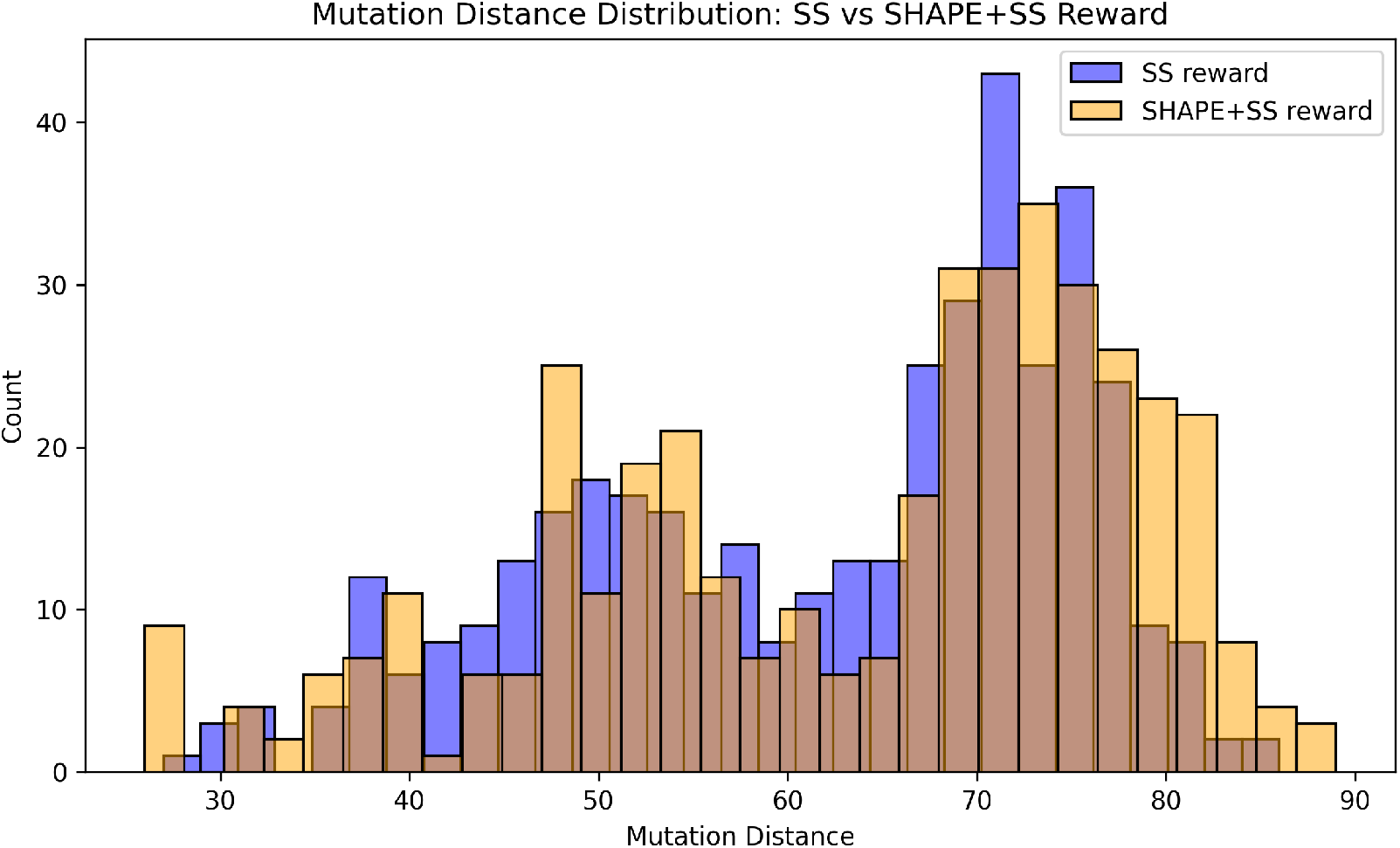
Distribution of mutation distances between generated sequences and wild-type sequences for Struct2SeQ and Struct2SeQ-SHAPE on OpenKnot 7 challenge.

Next we assess the experimental performance of Struct2SeQ designs in the OpenKnot Round 7 challenge, in comparison with other automated design methods and Eterna players. We note that our training and inference code at the time of this challenge had an error with OpenKnot score calculation, because a wrong threshold of SHAPE was used. It is possible that the wrong threshold caused significant performance degradation in terms of experimental OpenKnot scores, although we cannot quantify the exact impact of this error, due to the time limited nature of the challenge. Nevertheless, despite this error, Struct2SeQ-SHAPE designs still achieved human competitive experimental performance (p=0.29 compared to Eterna players by Wilcoxon two-sided test) that is competitive with Eterna players (humans) and other automated design methods (fig. 7). Interestingly, Struct2Seq performed much worse than Struct2SeQ-SHAPE in terms of experimental OpenKnot scores, despite computational metrics that are on par with Struct2SeQ-SHAPE. This suggests that the SHAPE constraints used in Struct2SeQ-SHAPE are critical for generating sequences that perform well experimentally, highlighting the importance of incorporating experimental constraints into the training process. Importantly, despite the error in OpenKnot score calculation, Struct2SeQ-SHAPE was still able to demonstrate human competitive performance, indicating the robustness of our approach.Overall, these results demonstrate the effectiveness of our reinforcement learning framework for RNA pseudoknot design, achieving human-level performance in a challenging design task while also generating chemically plausible sequences that perform well experimentally.

**Figure 7:**
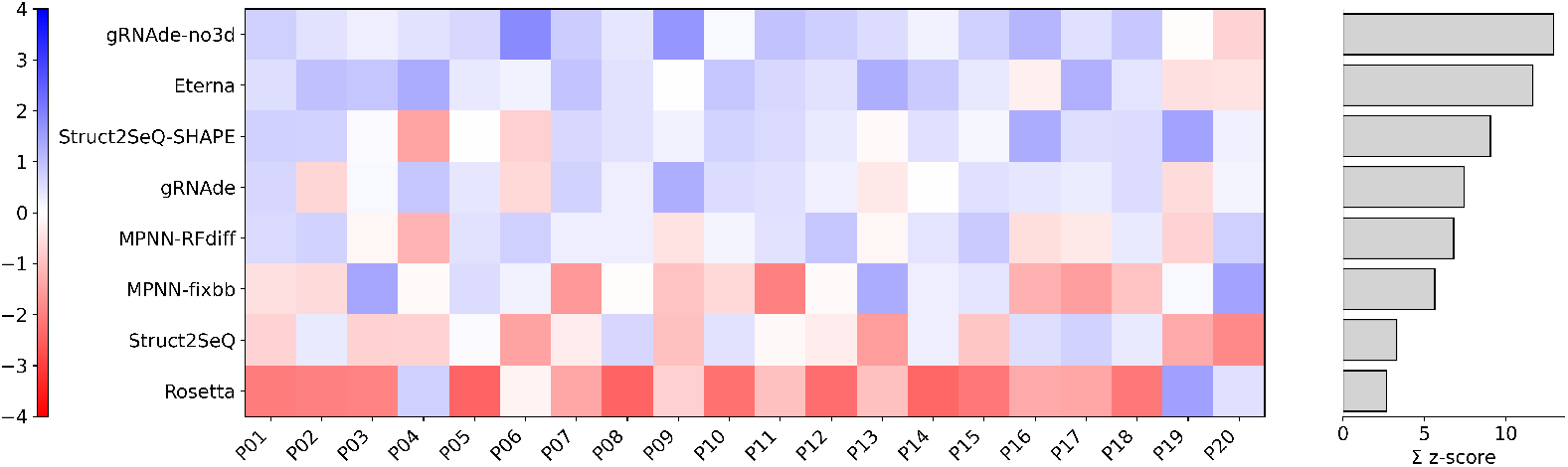
Struct2SeQ-SHAPE achieves human competitive performance in 100mer design challenge. Experimental results from OpenKnot Round 7 challenge. Here we show Z scores of 80th percentile OpenKnot scores for each method across all puzzles. Summed Z scores are from positive Z scores only following CASP RNA.

**Figure 8:**
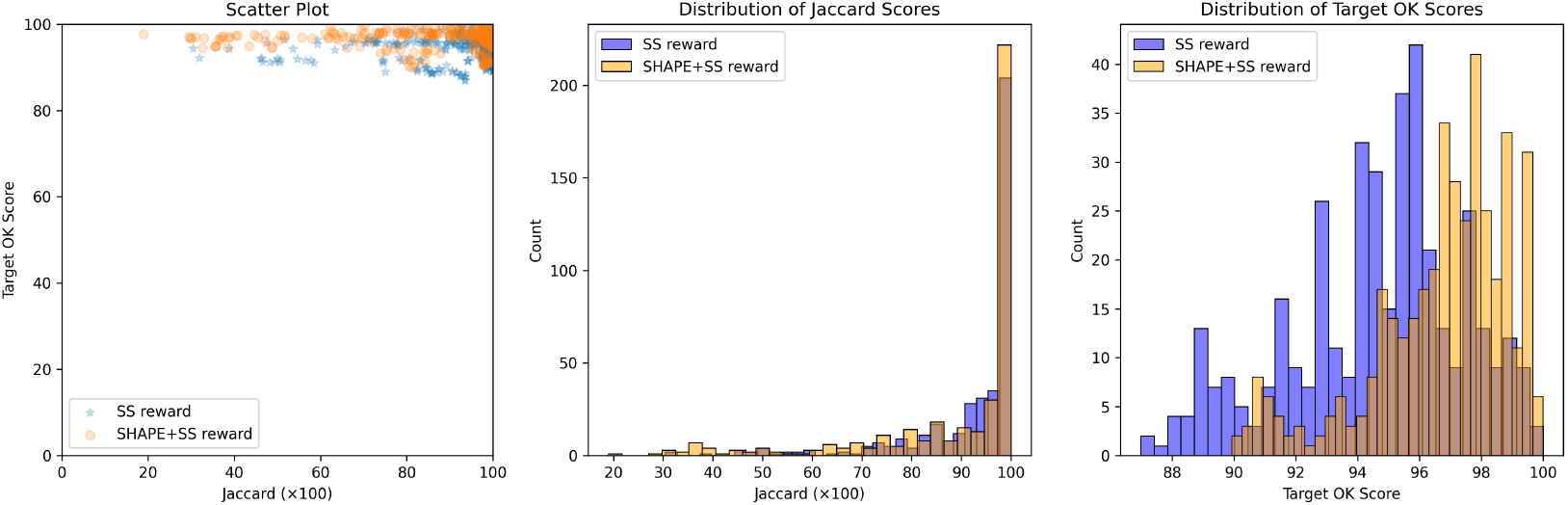
Performance of Struct2SeQ and Struct2SeQ-SHAPE on the OpenKnot 7b 240mer challenge. Left: Jaccard of base-pairs score for generated sequences vs scatter plot of Target OpenKnot score. Middle: histogram of Jaccard of base-pairs scores for generated sequences. Right: histogram of Target OpenKnot scores for generated sequences

### 3.2 OpenKnot Round 7b Pseudoknot design (240mer)

We also participated in the OpenKnot 7b 240mer challenge for designing RNA sequences that fold into pseudoknotted structures. To handle longer 240mer targets, we scaled Struct2SeQ and Struct2SeQ-SHAPE to 32 A100 nodes (256 GPUs) on 4000 A100 hr compute budget (i.e. on Nvidia DGX Clouds for 15 hrs on 32 nodes) and trained a new version of Struct2SeQ and Struct2SeQ-SHAPE on 240mer target structures from the human genomes as predicted by RibonanzaNet-SS. We scanned the human genome in 240mer windows with a stride of 50 and predicted secondary structures and SHAPE profiles using RibonanzaNet-SS and used the resulting structures with high OpenKnot scores as training target structures. Similar to 100mer designs in Openknot Round7, we set a compute budget where we generated 98000 sequences per target structure. While as expected, Struct2SeQ-SHAPE generated sequences with better OpenKnot scores compared to Struct2SeQ, Struct2SeQ-SHAPE matched Struct2SeQ in terms of Jaccard of base-pairs scores (fig. 8). This suggests that scaling to larger compute budgets may help Struct2SeQ-SHAPE match Struct2SeQ in terms of matching target secondary structures while still generating sequences with better OpenKnot scores. Similar to OK7 100mer challenge, we found that both Struct2SeQ and Struct2SeQ-SHAPE were able to generate sequences that match the target secondary structures at human-level performance while conforming to SHAPE constraints. interestingly, in some target structures, Struct2SeQ-SHAPE was able to generate more solutions that match the target secondary structure compared to Struct2SeQ (fig. 9 and fig. 10), suggesting that scaling Struct2SeQ-SHAPE to larger compute budgets may help it match or even outperform Struct2SeQ in terms of number of puzzles solved while still generating sequences with better OpenKnot scores.

**Figure 9:**
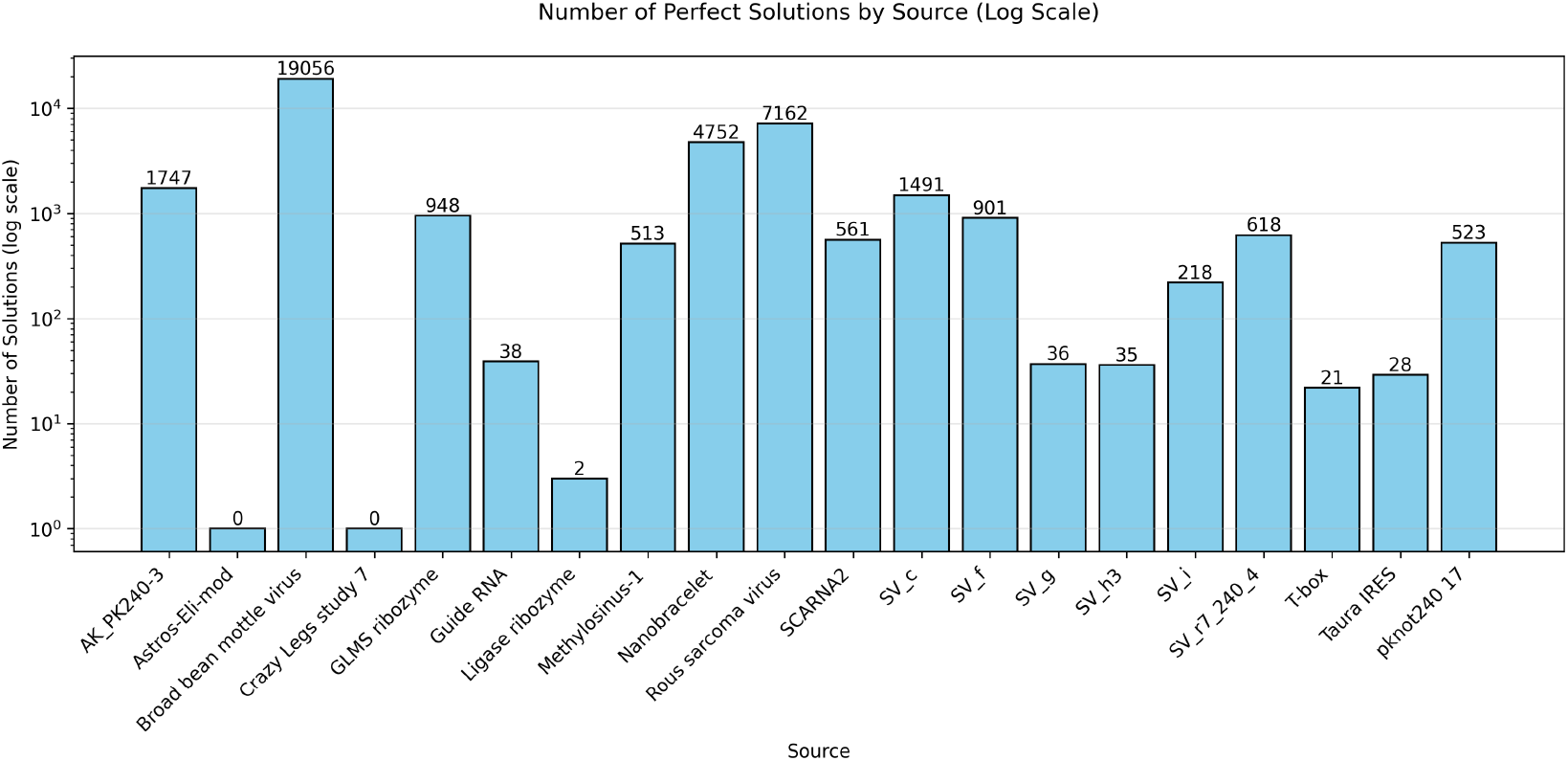
Number of solutions (matching base pairs) for each puzzle in the OpenKnot 7b 240mer challenge found by Struct2SeQ.

**Figure 10:**
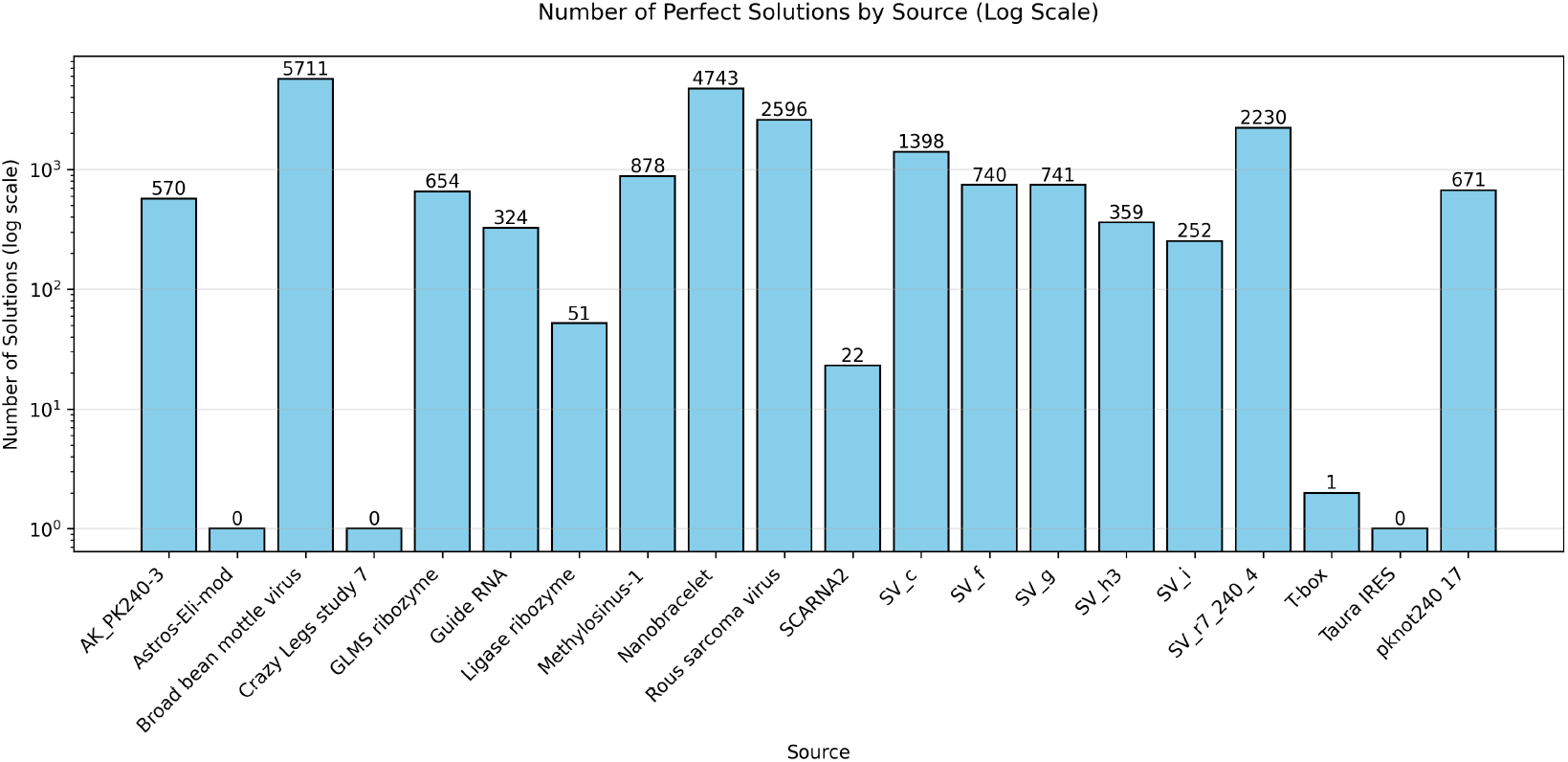
Number of solutions (matching base pairs) for each puzzle in the OpenKnot 7b 240mer challenge found by Struct2SeQ-SHAPE.

Again, we experimentally validate our designs in the OpenKnot 7b challenge. While here Struct2SeQ designs achieved human competitive performance (p=0.84 compared to Eterna players by Wilcoxon two-sided test), Struct2SeQ-SHAPE designs significantly outperformed Eterna players (p=0.024 compared to Eterna players by Wilcoxon two-sided test), with 2 times the summed z-scores compared to Eterna players (fig. 11). All other automated design methods are significantly outperformed by Struct2SeQ, Struct2SeQ-SHAPE, and Eterna players in this challenge. While gRNAde-no3d performed well in the OpenKnot 7 100mer challenge, it performed poorly in the OpenKnot 7b 240mer challenge. This is likely due to the increased sequence length and structural complexity. Further, other deep learning based design methods are all trained on sequence/structure pairs in the protein data bank, the majority of which are around 100nt in length, so these models likely do not generalize well to longer 240mer sequences. The success of Struct2SeQ and Struct2SeQ-SHAPE in this challenge also highlights that RibonanzaNet and RibonanzaNet-SS are able to accurately predict secondary structures and SHAPE profiles for longer 240mer sequences, which is critical for the success of our reinforcement learning framework. We also note that here we used the correct SHAPE thresholds during training and inference, which likely contributed to the strong experimental performance of Struct2SeQ-SHAPE designs. Overall, these results demonstrate the effectiveness of our reinforcement learning framework for RNA pseudoknot design, significantly outperforming human players and other automated design methods.

**Figure 11:**
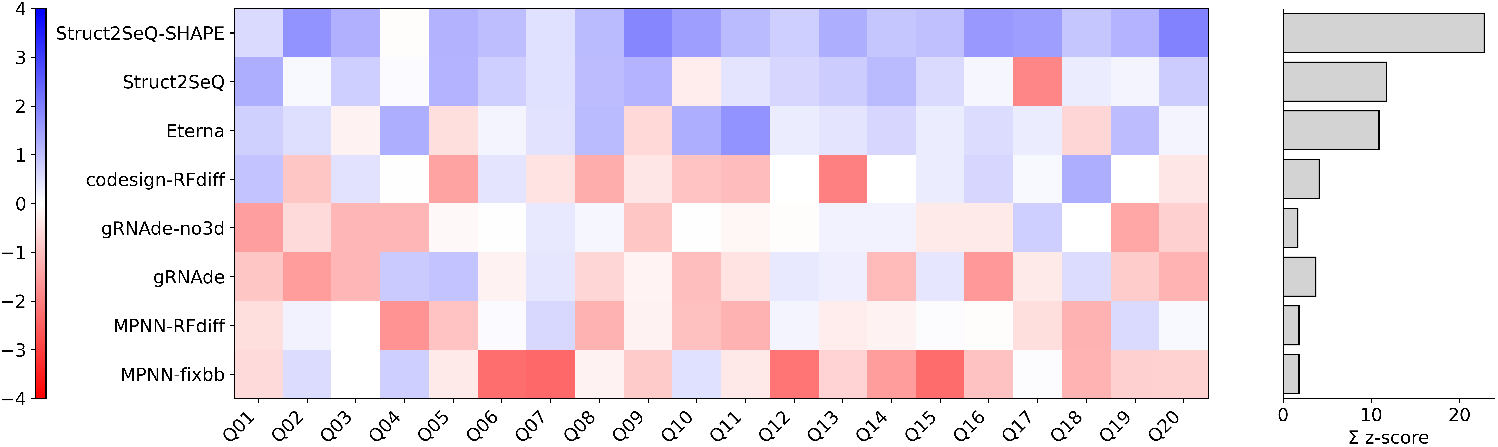
Struct2SeQ-SHAPE outperforms human and other automated methods in 240mer design challenge. Here we show Z scores of 80th percentile OpenKnot scores for each method across all puzzles. Summed Z scores are from positive Z scores only following CASP RNA.

## 4 Discussion

We have presented Struct2SeQ, a reinforcement learning framework for RNA inverse folding that integrates secondary structure and SHAPE reactivity constraints via deep Q-learning. By formulating RNA design as a sequential decision-making process, our model learns to generate sequences that not only fold into target secondary structures but also exhibit consistent SHAPE profiles. The use of position-wise rewards based on structural correctness and SHAPE compliance enables effective model training. Our results on the OpenKnot challenges demonstrate that Struct2SeQ achieves human-level performance in designing RNA sequences for complex pseudoknotted structures, while also generating diverse solutions that explore a broader sequence space compared to human players. The incorporation of SHAPE-informed rewards further enhances the chemical validity of generated sequences.

While in this work we focus on secondary structure design under SHAPE constraints, our reinforcement learning framework is in theory compatible with arbitrary reward functions, as long as they can be computed. This opens up exciting possibilities for other RNA design tasks, such as tertiary structure design, and design of RNAs with specific functional properties. Additionally, our results successfully scaling Struct2SeQ to 256 A100s suggest that scaling the model and training to larger compute budgets may unlock even better performance and solution diversity.

A limitation of the current reward design is that Struct2SeQ-SHAPE rewards are multiplicative combinations of structure correctness and SHAPE compliance, which means that if a position is incorrectly paired, it receives zero reward regardless of SHAPE compliance, so it might be helpful to decouple these two terms in future work. Specifically, we may ask the reinforcement learning agent to predict 2 separate Q-values, one for structure correctness and one for SHAPE compliance, and combine them in a weighted manner to compute the overall reward. This may help the model better learn to satisfy both structural and SHAPE constraints. Further, because the reinforcement learning agent is a neural network that can predict an arbitrary number of outputs, we may also consider predicting additional Q-values for other objective functions. For example, since RibonanzaNet does not consider thermodynamics, we may use EternaFold or ViennaRNA to compute the minimum free energy (MFE) structure of the generated sequence, and define an additional reward based on whether the MFE structure matches the target secondary structure. This may help generate sequences that are more thermodynamically stable in addition to satisfying structural and SHAPE constraints. Overall, our reinforcement learning framework provides a flexible and powerful approach for RNA design, and we believe it can be extended to tackle a wide range of RNA engineering challenges in the future.

Another limitation of the current work is the left to right decoding order. While all decoding orders are likely equivalent when averaged over all possible target structures, certain decoding orders may be more effective for specific structures. RNA design may also benefit from different decoding orders that better capture pairwise structural dependencies. For example, we may consider decoding paired positions together, or starting from loop regions and expanding outwards. Exploring alternative decoding strategies that better align with RNA folding principles may further improve design quality. To do this, we simply need to condition the model on the incomplete decoding sequence by randomly permuting the decoding sequence during training while applying the causal mask. During inference, arbitrary decoding orders can be applied.

## 5 Code Availability

Inference code for Struct2SeQ and Struct2SeQ-SHAPE is available at https://github.com/Shujun-He/Struct2SeQ.

## 6 Acknowledgements

We would like to thank Rhiju Das and the Eterna team for organizing the OpenKnot challenges and Jill Towney and Jonathan Romano for providing valuable target structures for training of 100mer Struct2SeQ models.

